# A digitization theory of the Weber–Fechner law

**DOI:** 10.1101/2021.03.15.435555

**Authors:** Haengjin Choe

## Abstract

Ever since the publication of Shannon’s article about information theory, there have been many attempts to apply information theory to the neuroscience field. Meanwhile, the Weber– Fechner law of psychophysics states that the magnitude of a subjective sensation of a person increases in proportion to the logarithm of the intensity of the external physical-stimulus. It is hardly surprising that we assign the amount of information to the response in the Weber– Fechner law. But, to date no one has succeeded in applying information theory directly to that law: the direct links between information theory and that response in the Weber–Fechner law have not yet been found. The proposed theory unveils a link between information theory and that response, and differs subtly from the field such as neural coding that involves complicated calculations and models. Because my theory targets the Weber–Fechner law which is a macroscopic phenomenon, this theory does not involve complicated calculations. My theory is expected to mark a new era in the fields of sensory perception research. My theory must be studied in parallel with the fields of microscopic scale such as neural coding. This article ultimately aims to provide the fundamental concepts and their applications so that a new field of research on stimuli and responses can be created.

## 1 Introduction

### 1.1 Psychophysics of Weber and Fechner

In 1834, the German physiologist Ernst Heinrich Weber conducted an exciting experiment. Weber gradually increased the weight which the blindfolded person was holding. With doing so, Weber asked the experimental subject when the increase of weight was first felt. Through this experiment, Weber found out that the least perceptible difference in weight was proportional to the starting value of weight. For instance, if someone barely feels the difference between 100 g and 104 g, then they barely feel the difference between 150 g and 156 g. At this time, a proportional expression 100:104=150:156 holds. Weber’s findings were elaborated in his book.^1^

Weber’s empirical observations were expressed mathematically by the German physicist Gustav Theodor Fechner who was Weber’s student.^2^ Fechner called his formulation Weber’s law. Fechner applied the result of Weber’s experiment to the measurement of sensation.^3,4^ The Weber–Fechner law established in the field of psychophysics attempts to mathematically describe the relationship between the magnitude of an external physical-stimulus and the perceived intensity of that stimulus.^2^ In this fascinating law, the physical stimuli are weight, sound frequency, etc. The Weber–Fechner law states that the relationship between physical stimulus and its perception is logarithmic. Therefore, if a stimulus varies as a geometric progression, the corresponding perception varies as an arithmetic progression. Bear in mind that the stimuli in the Weber–Fechner law have physical units but that the responses in that law have no physical units. The best example of the Weber–Fechner law is that many musicians adopt the equal temperament when they tune their musical instruments. Since the nineteenth century, musicians have tuned virtually all instruments in equal temperament.^5^ Equal temperament in music theory is a tuning system in which one octave is divided into 12 semitones of equal size; note that size here does not indicate frequency, but musical interval. The frequency corresponds to the stimulus, and the musical interval corresponds to the response. In the equal temperament system, the musical instruments are tuned so that the frequencies of all notes form a geometric sequence. Such a tuning results in the fact that the musical intervals of every pair of two consecutive notes are the same.^5,6^ This example needs to be explained in a bit more detail. Transposing or transposition in music means changing the key of a piece of music. When the musical-instrument accompaniment is too high or too low for us to sing to, we change the musical accompaniment into a lower or higher key. Even if we transpose the accompaniment into another key, as if by magic, we don’t feel awkward. There is an essential reason that the accompaniment does not sound awkward. This is because the musical instruments were tuned to the equal temperaments.

Such a tuning gives an equal perceived step size because pitch is perceived as the logarithm of sound frequency. As another concrete example of the Weber–Fechner law, response of the human ear varies logarithmically with sound intensity. Therefore, it is convenient to speak of the sound intensity level defined as

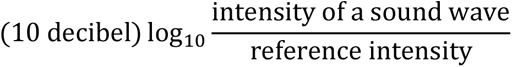

above the reference intensity.^7–16^ The reference intensity is chosen as 10^−12^ W/m^2^.^7–16^ This value is near the lowest limit of the human range of hearing at 1000 Hz.^7,10,12,13,15,16^ Note here that because the defining equation of the decibel scale has precisely the same form as the Weber–Fechner formula which we’ll soon see, the perceived magnitude is proportional naturally to the number obtained after we drop decibel in the sound level. The intensity level is a dimensionless quantity. Intensity level has a dimensionless unit associated with it, called the decibel. As a third concrete example of the Weber–Fechner law, response of the human eye varies logarithmically with brightness of light. The stellar magnitudes *m*_1_ and *m*_2_ for two stars are related to the corresponding brightnesses *b*_1_ and *b*_2_ through the relationship

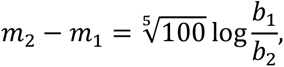

which results from the Pogson’s proposal that the difference of five magnitudes should be exactly defined as the brightness ratio of 100 to 1.^17–20^ Note here that the stellar magnitude corresponds to the response and hence has no physical unit. Stellar magnitude is dimensionless, since it’s formed from the ratio of two brightnesses.

### 1.2 Information theory of Shannon

Claude Elwood Shannon defined mathematically the concept of the ‘amount of information’ which is sometimes called the ‘information content’.^21^ Going one step further, he defined mathematically the concept of information entropy.^21^ Information entropy is similar to entropy in thermodynamics. Due to information theory, we came to be able to quantify the amount of information transmitted over a communication channel.

### 1.3 Digitization procedure

Is it possible to digitize the Weber–Fechner law? Digitizing the Weber–Fechner law in this article means the following: we define mathematically a new kind of the information content from the Weber–Fechner law, and going one step further we define mathematically a new kind of entropy from that law. This digitization will be able to be possible if we discover links between the information content formula of Shannon and the Weber–Fechner formula. I propose a theory that we are able to digitize the Weber–Fechner law. The digitization procedure is as follows. We will add one more stimulus to the existing various stimuli applied to the Weber–Fechner law. This addition procedure is based on the structural analogy between the Weber–Fechner formula and the information content formula. This additional stimulus and the corresponding perception are extremely extraordinary. We will digitize the Weber–Fechner law by extending and reinterpreting that law. The digitization of the Weber– Fechner law which is achieved by merging information theory with the Weber–Fechner law will lead us to two new concepts. Firstly, we define a new concept which I call the ‘amount of response information’ or the ‘response information content’. Ultimately, we end up defining another new concept which I call the ‘perception entropy’.

## 2. The Weber–Fechner law and the information content

### 2.1 Derivation of the Weber–Fechner law

The Weber–Fechner law is derived as follows.^22^ In Weber’s experiment, let *S* be weight at some instant and let d*R* be the differential increment in weight. Let d*R* be the differential change in perception. The equation to express the result of Weber’s experiment is

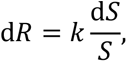

where *k* is a proportionality constant. Integrating both sides of this equation yields

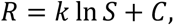

in which case *C* is an integration constant. Let us suppose that *S*_0_ is the threshold of the stimulus below which a person does not perceive anything. Because *R* = 0 whenever *S* = *S*_0_, the *C* must be equal to −*k* ln *S*_0_. Therefore, we can finally obtain

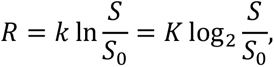

which is the mathematically expressed Weber–Fechner law. (There is one significant thing about *K*, which will be mentioned in Subsection 4.2.) Let us note that the *R* has no physical unit. But the physical stimulus *S* has, of course, a physical unit.

### 2.2 Definition of the information content

Shannon defined the amount of information of an event of the occurrence probability *P* as

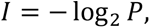

where the information content *I* is measured in bits.^21^ And he defined the entropy of a discrete random variate as the average amount of information of the random variate.^21^ In order to derive a new kind of the information content and hence a new kind of the entropy, let’s redescribe the above equation as

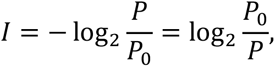

where as a matter of course *P*_0_ = 1.

## 3 Derivation of my theory

### 3.1 Special derivation

In the equation

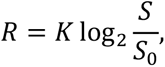

for all practical purposes *S* ≥ *S*_0_. In the equation

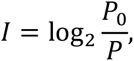

for all practical purposes *P* ≤ *P*_0_. We can interpret *P*_0_ = 1 as the threshold of *P*. Most importantly, we can interpret *I* as the response to *P*; it would be nice for us to call *P* a bizarre kind of stimulus. This bizarre stimulus fundamentally differs from other ordinary physical stimuli in that it is just a number. This stimulus has no physical unit. Very important another fact is that the higher *P* is, the lower *I* is but that the higher *S* is, the higher *R* is. But it is not essential that *I* is a strictly decreasing function with respect to *P* whereas *R* is a strictly increasing function with respect to *S*. The essentially important facts are that *R* and *I* with respect to *S* and *P*, respectively, are logarithmic and that both *S* and *P* have the threshold values. From these viewpoints, consequently, I cannot choose but say that the occurrence probability is an anomalous kind of stimulus. It would be appropriate that we name the occurrence probability the ‘mathematical stimulus’ in that the occurrence probability has no physical unit. This new type of stimulus is under the government of the Weber–Fechner law. By widening the range of applications of the Weber–Fechner law, we come to be capable of merging information theory with the Weber–Fechner law of psychophysics.

For simplicity, in current subsection we confine our attention to special situations: we set *K* to 1 for all sensations. (We make complete descriptions in Subsection 3.2. In that subsection we make no assumptions about *K*.) Then, we can say that the amount of perception is 0 bits when *S* = *S*_0_ and that the amount of perception is 1 bits when *S* = 2*S*_0_. However, there is one thing to be cautious about. In reality, different people have different *S*_0_’s. So, for the identical *S*, different people have different amounts of response information. In addition, as the body of a person ages, *S*_0_ is changed. So, for the identical *S* the response information content of even a person varies with age. Because of this fact, the response information content is quite different from the information content of Shannon. In the defining equation of the information content of Shannon, *P*_0_ is fixed at 1.

If the physical stimulus is the weight, the problem is simple. In order to clearly define a new kind of entropy, let’s consider the sound wave from now on. Because a pure tone which is a sound wave with sinusoidal waveform is characterized by both a frequency and a displacement amplitude, considering only either frequency or amplitude may be meaningless. Hence, we consider both simultaneously. For the sake of argument let’s consider a pure tone of frequency *f* and amplitude *A*. The Shannon entropy and physics entropy are subject to the constraint that the total sum of probabilities is 1. On the other hand, there is not such a constraint in the defining equation of the new kind of entropy. The physical quantities *A* and *f* change much more freely. Therefore, it is impossible that we define the new entropy in the same form as Shannon’s entropy. In order words, we must not define the new entropy as

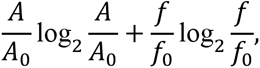

which is in the form of the familiar Shannon’s entropy. Before we define the new entropy, we first define the total amount of response as

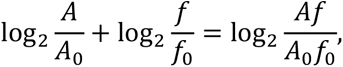

where 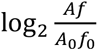 is measured in bits. In other words, the total amount of response is defined so that it equals the sum of the amounts of each individual response. Consequently, we define the new entropy as the arithmetic mean of the response information contents being considered; that is, new entropy is defined as

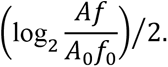

The unit of this quantity is bits/response. Please note that in the current situation there are two responses: one response due to the amplitude and the other due to the frequency. I named this new quantity the ‘perception entropy’ or the ‘percentropy’. The term ‘percentropy’ is a new compound word formed from perception and entropy. The percentropy does not correspond perfectly to the Shannon entropy. Nonetheless, it is useful to define the concept of percentropy. One of the reasons for this definition is that the percentropy is linked directly with the physics energy. We are easily able to understand such a fact. As an example, consider two pure tones. One tone has amplitude *A*_1_ and frequency *f*_1_. The other tone has amplitude *A*_2_ and frequency *f*_2_. The percentropy of the former tone is 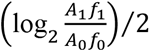. The percentropy of the latter tone is 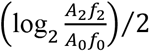. The *A*_1_ *f*_1_ = *A*_2_ *f*_2_ is a necessary and sufficient condition for the fact that the percentropies of the two tones are the same. And, *A*_1_ *f*_1_ = *A*_2_ *f*_2_ if and only if (*A*_1_ *f*_1_)^2^ = (*A*_2_ *f*_2_)^2^. We can therefore immediately conclude that the *A*_1_ *f*_1_ = *A*_2_ *f*_2_ is a necessary and sufficient condition for the fact that the energies of the two tones are the same. Let us now consider two compound tones. One compound tone consists of a pure tone which has amplitude *A*_11_ and frequency *f*_11_ and a pure tone which has amplitude *A*_12_ and frequency *f*_12_. The other compound tone consists of a pure tone of amplitude *A*_21_ and frequency *f*_21_ and a pure tone of amplitude *A*_22_ and frequency *f*_22_. Understandably, the total amount of response information of a compound tone is the sum of the total amounts of response information of constituent pure tones. In our situation, the total amount of response information of a compound tone is the sum of the total amount of response information of either constituent pure-tone and the total amount of response information of the other constituent pure-tone. The percentropy of the former compound-tone is

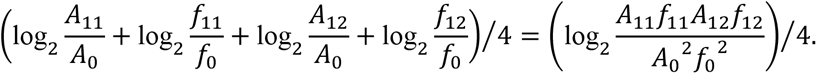

The percentropy of the latter compound-tone is

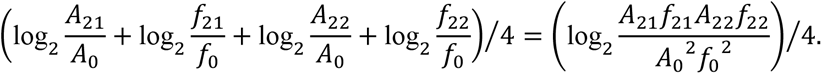

The *A*_11_*f*_11_*A*_12_*f*_12_ = *A*_21_*f*_21_*A*_22_*f*_22_ is a necessary and sufficient condition for the fact that the percentropies of the two compound-tones are the same. And, *A*_11_*f*_11_*A*_12_*f*_12_ = *A*_21_*f*_21_*A*_22_*f*_22_ if and only if (*A*_11_*f*_11_)^2^(*A*_12_*f*_12_)^2^ = (*A*_21_*f*_21_)^2^(*A*_22_*f*_22_)^2^. Therefore, in this compound-tones situation, the equality of the two percentropies is a necessary and sufficient condition for the fact that the products of energies of the two pure-tones composing each compound-tone are the same.

### 3.2 General derivation

In the last subsection, we conceptualized a few fundamental things in special situation—that is, when the *K*’s are 1 for all sensations. We now turn to the more general situation, in which the *K* has its own actual value for each sensation. Because the *K*’s are simply constants, we just have to define the response information content for a single stimulus as

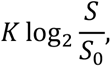

where 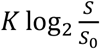 is measured in bits. Let’s consider a pure tone of frequency *f* and amplitude *A*. Let *K*_f_ and *K*_a_ be the constants corresponding to the frequency and the amplitude. We first define the total amount of response as

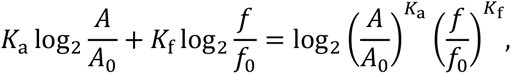

where 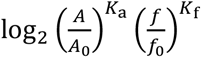 is measured in bits. In other words, the total amount of response is defined so that it equals the sum of the amounts of each individual response. And then we define the new entropy as the arithmetic mean of the response information contents being considered; that is, new entropy is defined as

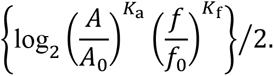

The unit of this quantity is bits/response. Because the amplitude and frequency change independently of each other, we took the arithmetic mean. Notice that the percentropy is not linked directly with the physics energy unless *K*_a_ equals *K*_f_.

## 4 Discussion

### 4.1 Utilization and interpretation

According to my theory, it comes to be possible that we utilize percentropy (or the total amount of response information) for quantifying the sensitivity of the sensory organs of humans; at this time the sensitivity is a function of stimuli and their thresholds. This quantification allows us to compare the sensitivities of the sensory organs of people. Amazingly, the sensitivity of the sensory organ implies the superiority of the sensory organ. It is best to explain a numerical example. We shall illustrate the calculation of the percentropy through a simple example. For simplicity, we set *K* = 1 for all sensations. Let us consider the case of the hearing. There are two people. For a person X, *A*_0_ = 33 pm and *f*_0_ = 20 Hz. For another person Y, *A*_0_ = 22 pm and *f*_0_ = 60 Hz. Suppose, in addition, that the two people hear a sound wave with an amplitude of 330 pm and a frequency of 1000 Hz. Then, the percentropy in the hearing organ of X is

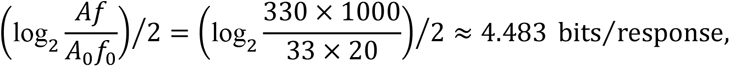

and that of Y is

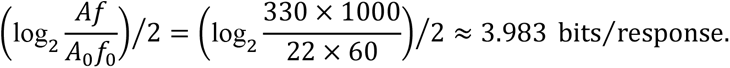

Therefore, the hearing organ of X is superior to that of Y. In fact, as we can very easily understand, we only need to know each individual’s *A*_0_*f*_0_ in comparing superiority of the hearing organs of people. Consequently, we can say that the information-theoretical measure of the superiority of the hearing organs of people is *A*_0_*f*_0_. Of course, we might effortlessly devise the *A*_0_*f*_0_ as a measure of the superiority, without any knowledge appearing in this article. The key point here is that the measure *A*_0_*f*_0_ is justified by the digitization theory of the Weber–Fechner law; i.e., the propostion that *A*_0_*f*_0_ is a measure of the superiority is valid from the information-theoretical perspective. Please note that we have set *K* = 1 regardless of the type of sensation.

What is the appropriate generalization of *A*_0_*f*_0_? Even if we do not assume that *K* = 1 for all sensations, we can still compare the sensitivity of the sensory organs of people. But the measure of the superiority is no longer *A*_0_*f*_0_. If we assume that the values of *K*_a_’s for people are all identical and that the values of *K*_f_’s for people are all identical, then the information-theoretical measure of the superiority is 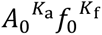.

The example of current subsection validates my interpretation of the Weber–Fechner formula as defining the response information content. Okay. Things are going well so far. The question is then what 4.483 bits/response and 3.983 bits/response mean; equivalently, what 8.966 bits and 7.966 bits mean. Suppose constant physical-stimulus is exerted on people. It is clear that (1) there is no significant difference between the duration of the external stimulus and the duration of the response, and (2) the response information content is proportional to the duration of the response. And, presumably, the duration of the response is quantized. In order words, the duration of the response is always some integral multiple of elementary duration. According to this hypothesis, the response information content during a time interval equals some integer times the response information content during elementary duration. Consequently, while talking about the numerical value of *K*, we must mention the duration of the response together. However, it is cumbersome to do so. Therefore, it is convenient to determine the numerical value of *K* for elementary duration first and then use the value so determined. Now, we can say what 8.966 bits and 7.966 bits mean. The 7.966 bits is a certain number times the response information content for elementary duration.

### 4.2 Future challenges

The stimuli and responses that have been mentioned so far are about the external physical-stimuli and their perceptions; the stimuli and responses that have been mentioned so far do not include the stimuli and responses that neural coding handles. In order for the digitization theory of the Weber–Fechner law to be firmly established, we need to elucidate the relationships between microscopic and macroscopic theories on stimuli and responses.

Past information-theoretical researches on stimuli and responses have mainly focused on microanalysis.^23–30^ Those researches are accompanied by complicated models and calculations. In addition to microscopic approaches to sensory perception research, we need to take macroscopic approaches as well. Because the Weber–Fechner law is a macroscopic phenomenon which is a manifestation of numerous microscopic outcomes, the calculations involved in my theory are far less complicated. In the future, macroanalyses as well as microanalyses should be done together.

There is one significant thing about *K* in particular. In psychophysics *K* is originally an experimentally determined constant. But, the numerical value of *K* for each sense in my theory must be determined by comparing my theory with works of microscopic scale.

Meanwhile, I presume that regardless of the type of sense, a theoretic expression (not a measured value) for *K* includes the Boltzmann constant or Boltzmann-like constant which serves to bridge the macroworld and the microworld. Once the *K*’s and elementary durations are all determined, the field of research on stimuli and responses is expected to develop rapidly.

## 5 Conclusions

Whereas statistical mechanics is the most detailed level for the treatment of phenomena in systems with many particles, thermodynamics discards the detailed dynamical evolution of a system.^31^ These two branches of physics complement each other. If we can say that neural coding corresponds to statistical mechanics, what can be said to correspond to thermodynamics? To our regret, there is nothing that can be said to correspond to thermodynamics. This reality motivates us to make a theory. The Weber–Fechner law is a macroscopic manifestation. Therefore, our new theory should be aimed at that law. So as to achieve our goal, we must find links between the information content formula and the Weber–Fechner formula. These links are the key idea behind the digitization theory of the Weber–Fechner law. The link presented in this article is an undiscovered link. Recognizing that we can redescribe the defining equation of the information content using the threshold of probability enabled the discovery of that link. After finding the link, we strictly conceptualized several things. The Weber–Fechner formula is similar in structure to the information content formula, and this similarity invites an analogous interpretation. Consequently, the definition of the information content for physiological response was chosen to be in analogy with the definition of the information content which had been given by Shannon; the unit of the response information content is bits. Based on the analogy between the defining equation of the information content and the formula describing the Weber–Fechner law, I was able to theorize about the digitization of this law. On the one hand, the response information content has some features of conventional information theory. On the other hand, the response information content has the different features from conventional information theory. We also defined the perception entropy, which is abbreviated as percentropy. The percentropy means the average amount of the human response to external physical-stimuli; the unit of the percentropy is bits/response. Percentropy is also very information-theoretic, in the sense that its definition originates in the merging of information theory with the Weber–Fechner law. Amazingly, we could judge the superiority of sensory organs of humans by percentropy. The percentropy affords a measure of how good a sensory organ is: as a sensory organ is more superior, the percentropy is greater. Because of such judgment, my theory is expected to find applications in medical diagnosis. The specific numerical value of percentropy is a new criterion for judging whether a certain part of an organism is normal or abnormal.

Neural coding treats the microworld such as neurons, so calculations in neural coding are complicated. My theory targets the Weber–Fechner law which is a macroscopic phenomenon, so calculations in my theory are simple. In order for my theory to be firmly established, we need to elucidate the relationships between microscopic and macroscopic theories on stimuli and responses. These relationships can be found by determining the values of the *K*’s and elementary durations. If my theory turns out to be of great usefulness, a new field of research will be created. Then, that new field will find practical applications in many diverse areas such as physiology and medicine. It will be good for us to name this new field ‘perceptional information theory’ or ‘perceptual information theory’.

